# Evaluation of the geographical utility of Eastern Russell’s viper (*Daboia siamensis*) antivenom from Thailand and an assessment of its protective effects against venom-induced nephrotoxicity

**DOI:** 10.1101/591305

**Authors:** Janeyuth Chaisakul, Nattapon Sookprasert, Robert A. Harrison, Narongsak Chaiyabutr, Lawan Chanhome, Nicholas R. Casewell

**Author notes:** **Correspondence to:**, Department of Pharmacology, Phramongkutklao College of Medicine, Bangkok 10400, Thailand.

## Abstract

**Background:** *Daboia siamensis* (Eastern Russell’s viper) is a medically important snake species found widely distributed across Southeast Asia. Envenomings by this species can result in systemic coagulopathy, local tissue injury and/or renal failure. While administration of specific antivenom is an effective treatment for Russell’s viper envenomings, the availability of, and access to, geographically-appropriate antivenom remains problematic in many rural areas. In this study, we determined the binding and neutralizing capability of antivenoms manufactured by the Thai Red Cross in Thailand against *D. siamensis* venoms from three geographical locales: Myanmar, Taiwan and Thailand.

**Methodology/ Principle findings:** The *D. siamensis* monovalent antivenom displayed extensive recognition and binding to proteins found in *D. siamensis* venom, irrespective of the geographical origin of those venoms. Similar immunological characteristics were observed with the Hemato Polyvalent antivenom, which also uses *D. siamensis* venom as an immunogen, but binding levels were dramatically reduced when using comparator monovalent antivenoms manufactured against different snake species. A similar pattern was observed when investigating neutralization of coagulopathy, with the procoagulant action of all three geographical venom variants neutralized by both the *D. siamensis* monovalent and the Hemato Polyvalent antivenoms, while the comparator monovalent antivenoms were ineffective. Assessments of *in vivo* nephrotoxicity revealed that *D. siamensis* venom (700 µg/kg) significantly increased plasma creatinine and blood urea nitrogen levels in anaesthetised rats. The intravenous administration of *D. siamensis* monovalent antivenom at three times higher than the recommended scaled therapeutic dose, prior to and 1 h after the injection of venom, resulted in reduced levels of markers of nephrotoxicity, although lower doses had no therapeutic effect.

**Conclusions/Significance:** This study highlights the potential broad geographical utility of the Thai *D. siamensis* monovalent antivenom for treating envenomings by the Eastern Russell’s viper. However, only the early delivery of high antivenom doses appear capable of preventing venom-induced nephrotoxicity.

**Author summary:** Snakebite is a major public health concern in rural regions of the tropics. The Eastern Russell’s viper (*Daboia siamensis*) is a medically important venomous snake species that is widely distributed in Southeast Asia and Southern China, including Taiwan. Envenoming by *D. siamensis* causes several systemic pathologies, most notably acute kidney failure and coagulopathy. The administration of antivenom is the mainstay therapeutic for treating snakebite, but in remote areas of Myanmar and Southern China access to antivenom is limited, and can result in the use of inappropriate, non-specific, antivenoms and treatment failure. Therefore, maximizing the utility of available efficacious antivenom is highly desirable. In this study, we investigated the utility of the widely available Thai Red Cross antivenoms for binding to and neutralizing *D. siamensis* venoms sourced from three distinct locales in Asia. Since the effectiveness and antivenom dose required to prevent *D. siamensis* venom-induced nephrotoxicity has been controversial, we also examined the preclinical efficacy of *D. siamensis* antivenom at preventing this pathology in experimentally envenomed anaesthetised animals. Our findings suggest that monovalent antivenom from Thailand, which is clinically effective in this country, has highly comparable levels of immunological binding and *in vitro* neutralization to *D. siamensis* venoms from Taiwan and Myanmar. We also show that the early administration of high therapeutic doses of antivenom are likely required to neutralize nephrotoxins and thus prevent acute renal failure following envenoming. Our findings suggest that certain Thai Red Cross antivenoms likely have wide geographical utility against *D. siamensis* venom and therefore may be useful tools for managing snakebite envenomings by this species in the absence of available locally manufactured therapeutics.

## 1. Introduction

Snake envenoming is one of the world’s most lethal neglected tropical diseases, resulting in as many as 138,000 deaths per year [1]. Snakebite predominately affects the rural poor populations of the tropics, with the regions of sub-Saharan Africa, South Asia and Southeast Asia suffering the greatest burden of both morbidity and mortality [1, 2]. One of the most medically important groups of venomous snakes are the Russell’s vipers (Viperidae: *Daboia* spp.). These relatively large, predominately nocturnal, snakes have a wide distribution across much of southern Asia [3] and have been classified into two species, the Western Russell’s viper (*Daboia russelii*) and the Eastern Russell’s viper (*Daboia siamensis*). The Western Russell’s viper is found distributed across India, Pakistan, Bangladesh and Sri Lanka, whilst the Eastern Russell’s viper has a wide distribution throughout Southeast Asia and Southern China, including Taiwan [4, 5] (Figure 1). In 2016, the World Health Organization (WHO) categorized *D. siamensis* as a snake causing high levels of morbidity and mortality in Myanmar, Thailand and some Indonesian islands, *i.e.* Java, Komodo, Flores and Lomblen [6].

**Figure 1.**
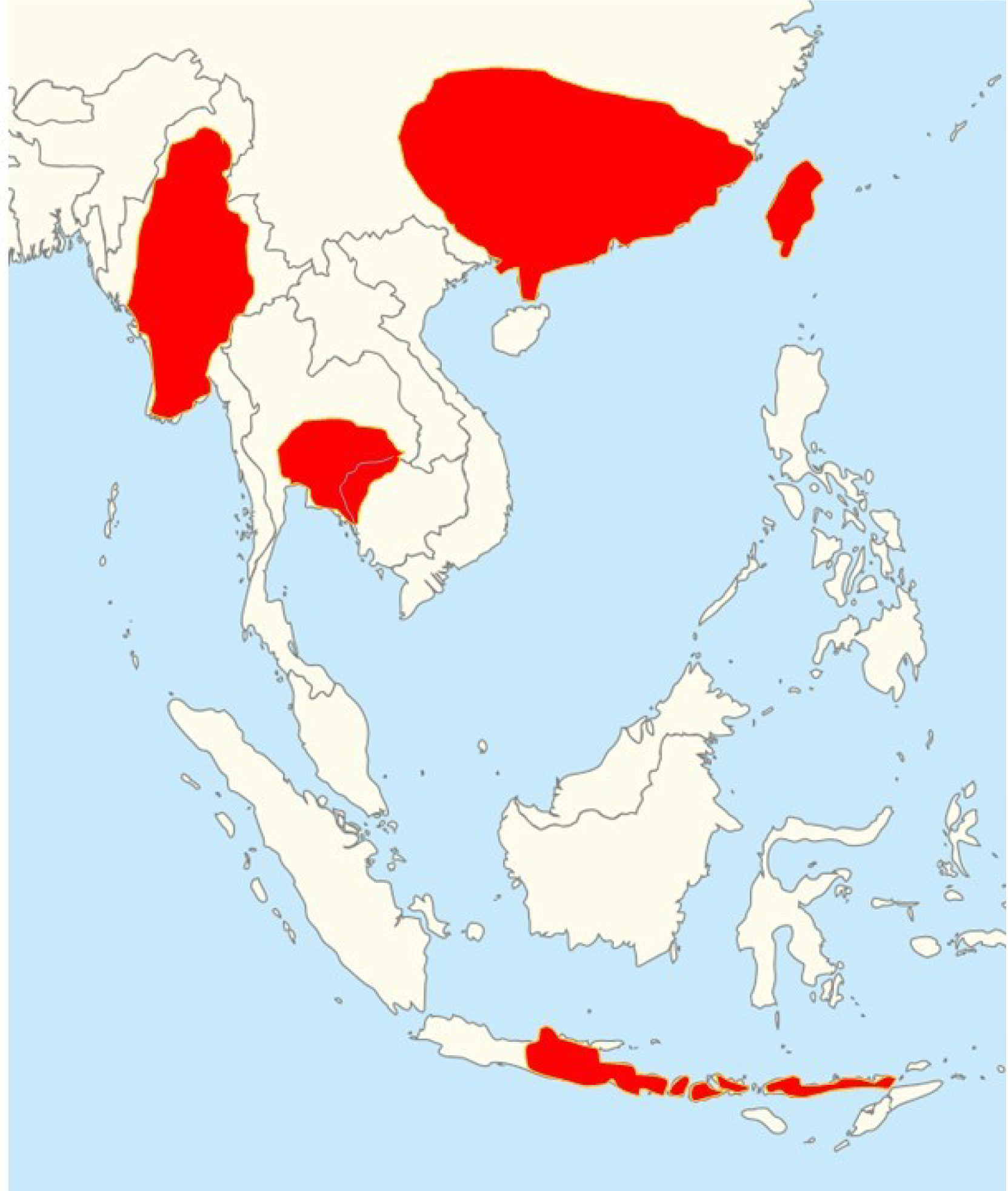
The distribution of *Daboia siamensis* in Asia [4, 5].

Envenomings by medically important Asian vipers are typically clinically characterized by hemodynamic alterations. Clinical outcomes following *D. siamensis* envenoming can include: local painful swelling at the bite-site, conjunctival oedema, systemic coagulopathy and/or haemorrhage, while hypopituitarism has also been reported [7]. In addition, *D. siamensis* venom can induce renal toxic effects (nephrotoxicity), which are characterized by hematuria, tubular necrosis and acute renal failure [8]. These variable clinical signs observed following snakebites are a consequence of Russell’s viper venoms exhibiting considerable variation across their range, resulting in differences in their toxin profiles, which in turn impacts upon clinical outcomes observed in snakebite victims [9].

Two major snake venom toxin families are thought to be predominately responsible for the bleeding disorders and renal failure observed following systemic envenoming by *D. siamensis*, the enzymatic phospholipases A_2_ (PLA_2_) and snake venom metalloproteinases (SVMP). Both toxin types are often found to be major components of viper venoms [10], but each toxin family is known to encode multiple isoforms that vary among species and are capable of exhibiting distinct functional activities [10, 11]. Such protein neo-/sub-functionalization is thought to be underpinned by multiple gene duplication events coupled to accelerated bursts of adaptive evolution [12–14]. Consequently, snake venom PLA_2_s are responsible for several pharmacological activities including neurotoxicity, myotoxicity, anticoagulant effects, smooth muscle relaxation/hypotension and hypersensitivity [15], while SVMP functional activities include the activation of different coagulation factors and the degradation of endothelial cell membranes, resulting in venom-induced consumption coagulopathy and haemorrhage [16]. Other toxin families (e.g. L-amino acid oxidases, serine proteases and C-type-lectin-like proteins [9]) likely contribute to pathology following envenoming by Russell’s vipers, and together with the PLA_2_ and SVMPs, these toxin families comprise >90% of all of the toxins found in the venom proteome [17]. In terms of specific toxins, prior studies have demonstrated that the SVMP RVV-X is a potent activator of Factor X [18], and thus contributes to the depletion of coagulation factors (notably fibrinogen), resulting in a syndrome similar to disseminated intravascular coagulation (DIC) termed venom-induced consumption coagulopathy (VICC) [19]. Moreover, both PLA_2_ and SVMP from *D. siamensis* venom have been demonstrated to initiate kidney injury via an increase in renal vascular resistance or renal ischemia, resulting in decreases in renal blood flow, glomerular filtration rate and urine flow [20, 21]. Finally, *D. siamensis* venom fractions enriched in PLA_2_ and SVMP toxins have been demonstrated to cause marked decreases in mean arterial pressure and also promoted the release of inflammatory mediators in aneasthetised dogs [20].

The only effective treatment for systemic snakebite envenoming is specific antivenom, which consists of polyclonal antibodies isolated from hyperimmune animal serum/plasma. Non-pharmacological treatments, which include the local use of tourniquets, cross-shaped skin incision, local suction or irrigation, or the administration of non-specific snake antivenom (*i.e.* made against snake species other than that which bit the patient) are typically ineffective, and potentially harmful [22]. There are two types of specific antivenom available for treating *D. siamensis* envenomings; monovalent (or monospecific) antivenom, which comprises of polyclonal antibodies derived from equine plasma hyperimmunized with *D. siamensis* venom only, and polyvalent (or polyspecific) antivenom, which consists of antibodies sourced from animals immunized with *D. siamensis* venom and venoms from other medically important snake species.

An example of this latter type of antivenom is made by The Queen Saovabha Memorial Institute (QSMI) of the Thai Red Cross Society in Bangkok, Thailand, which produces the Hemato Polyvalent Snake antivenom (HPAV) for treating viper envenomings from *D. siamensis, Calloselasma rhodostoma* and *Trimeresurus albolabris.* QSMI also produce monovalent antivenoms for *D. siamensis* (DSAV), C. *rhodostoma* (CRAV) and *T. albolabris* (TAAV), while other monospecific products against *D. siamensis* are manufactured elsewhere in Asia, such as Myanmar (Myanmar Pharmaceutical Factory) and Taiwan (Centre for Disease Control). Previous studies have reported a degree of cross-neutralizing effect between the HPAV and DSAV antivenoms from Thailand against the toxicity of *D. siamensis* venoms from different geographical localities, *i.e*. Myanmar and Taiwan, in preclinical studies and ELISA binding experiments, respectively, although their efficacy was seemingly lower than that of local antivenoms made using venom from those localities [17, 23]. However, despite the Myanmar Pharmaceutical Factory producing 46,000 vials of Russell’s viper antivenom annually, this volume is seemingly insufficient to treat the total burden of *D. siamensis* envenomings in country [24]. This therapeutic shortfall places an onus on the scientific community to robustly assess the likely therapeutic value of alternative antivenoms for treating *D. siamensis* envenomings in countries where antivenom supply may be restricted.

In addition, while antivenom remains the primary treatment for Russell’s viper envenoming across Asia, there has been considerable debate regarding its clinical effectiveness against venom-induced nephrotoxicity. In part, this stems from questions relating to the most appropriate dosing regimen for antivenom, and a lack of robust clinical studies relating to this topic. For example, even the use of high doses of monospecific antivenom (> 4 vials; 40 ml) in envenomed patients has been said to result in limited success in reversing progressive renal failure [7]. Consequently, dialysis (either peritoneal dialysis or haemodialysis) is often relied upon to manage such severe clinical outcomes. Despite the value of preclinical models for exploring antivenom efficacy, little research has been undertaken on the therapeutic value of antivenom at treating *D. siamensis*-induced nephrotoxicity. Leong *et al*. (2014) demonstrated that the Hemato Polyvalent antivenom (200 µl) exerted a protective effect on the occurrence of hematuria and proteinuria following the injection of *D. siamensis* venom within 4 h. However, due to the restricted monitoring time employed in this study, key markers of nephrotoxicity, such as blood urea nitrogen (BUN) and creatinine, were not detected [23].

In this study, we sought to further investigate the likely efficacy of antivenom against *D. siamensis* venoms sourced from different geographical locales. We used a variety of *in vitro* immunological and functional assays to assess the binding and neutralising effect of Thai (QSMI) antivenoms against *D. siamensis* venoms sourced from Thailand, Myanmar and Taiwan. Subsequently, we investigated the protective effect of the monospecific *D. siamensis* antivenom (DSAV) against the nephrotoxicity caused by *D. siamensis* venom *in vivo*, by quantifying plasma blood urea nitrogen (BUN) and creatinine levels in experimentally envenomed rats. Our findings demonstrate extensive antivenom cross-reactivity among geographical variants of *D. siamensis*, but that nephrotoxicity caused by *D. siamensis* venom is only inhibited when antivenom is delivered early and in high volumes. These results strongly advocate for further clinical research to be undertaken to validate the efficacy of *D. siamensis* antivenom across South-East Asia, particularly in systemically envenomed snakebite victims suffering from nephrotoxicity.

## 2. Materials and Methods

### 2.1 Snake venoms

Specimens of Thai Russell’s viper (*D. siamensis*) were maintained in captivity at QSMI, The Thai Red Cross Society Bangkok, Thailand. Venom was extracted from several snakes, pooled, and then frozen before being freeze-dried. Lyophilized *D. siamensis* venoms from Myanmar and Taiwan were provided from the Centre for Snakebite Research & Interventions, Liverpool School of Tropical Medicine, historical venom archive. Freeze-dried venom samples were stored at 4 °C, prior to use. Venoms were weighed, reconstituted in phosphate-buffered saline (PBS) and venom protein concentrations measured using a Nanodrop (ThermoFisher) and BCA protein assay (Pierce Biotechnology, Rockford, IL, USA).

### 2.2 Antivenoms

Hemato Polyvalent Snake antivenom (HPAV; Lot NO: HP00218, expiry date 03/2023), monovalent antivenoms for *D. siamensis* (DSAV; Lot NO: WR00117, expiry date 11/2022), *C. rhodostoma* (CRAV; Lot NO: CR00316, expiry date 06/2021) and *T. albolabris* (TAAV; Lot NO: TA00317, expiry date 07/2022) were purchased from QSMI of Thai Red Cross Society, Bangkok, Thailand. The freeze-dried antivenoms were dissolved with pharmaceutical grade water supplied by the manufacturer. The dissolved antivenoms were then stored at 4 °C prior to use. The protein concentrations were measured using a Nanodrop (ThermoFisher) and BCA protein assay (Pierce Biotechnology, Rockford, IL, USA). Normal horse IgG (1 mg/mL; Sigma, UK) was used as negative control.

### 2.3 Immunological assays

The protein concentrations of antivenoms were adjusted to 50 mg/ml for all immunological assays (original concentrations of reconstituted antivenoms were: HPAV, 54 mg/ml; DSAV, 21 mg/ml; CRAV, 24.5 mg/ml; and TAAV, 14 mg/ml).

#### 2.3.1 End Point Titration (EPT) ELISA

Immunological binding activity between venoms and antivenoms were determined following a previously described method [25]. First, 96 well ELISA plates (Nunc) were coated with 100 ng of venom (a separate plate for each Russell’s viper venom sample) prepared in carbonate buffer, pH 9.6 and the plates incubated at 4 °C overnight. Plates were washed after each stage using six changes of TBST (0.01 M Tris-HCl, pH 8.5; 0.15M NaCl; 1% Tween 20). Next, the plate was incubated at room temperature for 3 hours with 5% non-fat milk (diluted with TBST) to ‘block’ non-specific reactivity. The plates were then washed and incubated (in duplicate) with DSAV, CRAV, TAAV or HPAV antivenoms, at an initial dilution of 1:100 followed by 1:5 serial dilutions across the plate, and then incubated overnight at 4 °C. The plates were then washed again and incubated with horseradish peroxidase-conjugated rabbit anti-horse IgG (1:1000; Sigma, UK) for 3 hours at room temperature. The results were visualized by addition of substrate (0.2% 2,2/-azino-bis (2-ethylbenzthiazoline-6-sulphonic acid) in citrate buffer, pH 4.0 containing 0.015% hydrogen peroxide (Sigma, UK), and optical density (OD) measured at 405 nm. The titre is described as the dilution at which absorbance was greater than of the negative control (IgG from non-immunised horses; Sigma, UK) plus two standard deviations.

#### 2.3.2 Relative avidity ELISA

The chaotropic ELISA assay was performed as previously described [26]. In brief, the assay was performed as per the EPT ELISA assay detailed above, except that the antivenoms and normal horse IgG were diluted to a single concentration of 1:10,000, incubated overnight at 4 °C, washed with TBST and then a chaotrope, ammonium thiocyanate (NH_4_SCN), added to the wells in a range of concentrations (0-8 M) for 15 minutes. Plates were then washed again with TBST, and all subsequent steps were the same as the EPT ELISA. The relative avidity was determined as the percentage reduction in ELISA OD reading (measured at 405 nm) between the maximum (8 M) and minimum (0 M) concentration of NH_4_SCN.

#### 2.3.3 SDS-PAGE and immunoblotting

Lyophilized *D. siamensis* venoms were reconstituted to 1 mg/ml in reducing protein loading buffer and heated at 100 °C for five minutes. Nine µg of venom, together with molecular weight marker (Broad range molecular weight protein markers, Promega) was added to a 15% SDS-PAGE gel and separated under 200 volts, with the resultant proteins visualised by staining with Coomassie Blue R-250.

For immunoblotting, we repeated the electrophoresis experiments, except the gels were not stained, and were instead electro-blotted onto 0.45 µm nitrocellulose membranes using the manufacturer’s protocols (Bio-Rad, UK). Following confirmation of successful protein transfer by reversible Ponceau S staining, the membranes were incubated overnight in blocking buffer (5% non-fat milk in TBST), followed by six washes of TBST over 30 minutes and incubation overnight with primary antibody (*i.e.* the four antivenoms; HPAV, DSAV, CRAV, TAAV and horse IgG) diluted 1:5,000 in blocking buffer. Blots were washed as above, then incubated for 2 hours with secondary antibody *-* horseradish peroxidase-conjugated rabbit anti-horse IgG (Sigma, UK) diluted 1:1,500 with PBS. Then the membrane was washed again with TBST and visualised after the addition of DAB substrate (50 mg 3,3-diaminobenzidine, 100 ml PBS and 0.024% hydrogen peroxide; Sigma, UK).

### 2.4 In vitro coagulopathic activity

The neutralising effect of Thai antivenoms on the coagulopathic activity of *D. siamensis* venoms was determined using a previously described citrated bovine plasma coagulation assay [27]. Briefly, frozen bovine plasma (VWR International, Leicestershire, UK) was defrosted at 37 °C and centrifuged to remove precipitates (20-30 s at 1400 rpm). PBS (10 µL/well) was used as a control (PBS alone) as well as a diluent. Stock solutions of venom (100 ng/10 µL) were manually pipetted in triplicate into the wells of a 384 well microtiter plate. The wells were then overlaid with CaCl_2_; 20 mM (20 µL) and plasma (20 µL) using a Thermo Scientific Multidrop 384-autopipettor. To determine the protective effect of antivenom on clotting activity, we scaled therapeutic doses recommended by the manufacturer to the venom dose used as challenge (*i.e.,* 1 mL of HPAV and DSAV per 0.6 mg of venom, 1 mL CRAV per 1.6 mg venom, and 1 mL TAAV per 0.7 mg venom). To prepare the mixture, either HPAV (0.17 µL or 9.2 µg/well), DSAV (0.17 µL or 3.6 µg/well), CRAV (0.07 µL or 1.7 µg/well) or TAAV (0.15 µL or 3.6 µg/well) was mixed to the venom/PBS solution for 10 min prior to the addition of CaCl_2_; 20 mM (20 µL) and plasma (20 µL).

For all samples, we measured the kinetic absorbance at 25 ^°^C every 76 s for 100 cycles using a BMG Fluorostar Omega plate reader at 595 nm (BMG LABTECH, UK). Three different sources of data, consisting of single reading, a reading range, and average rate in time per well, were obtained for the determination of coagulation curves. In addition, the area under the curve (AUC) of each reaction was calculated and normalized as the percentage of venom clotting activity.

### 2.5 In vivo measures of nephrotoxicity

#### 2.5.1 Animal ethics and care

Male Sprague-Dawley rats were purchased from Nomura-Siam International Co. Ltd., Bangkok, Thailand. Rats were housed in stainless steel containers with access to food and drinking water *ad libitum*. Approvals for all experimental procedures were obtained from the Subcommittee for Multidisciplinary Laboratory and Animal Usage of Phramongkutklao College of Medicine and the Institutional Review Board, Royal Thai Army Department, Bangkok, Thailand (Documentary Proof of Ethical Clearance no: IRBRTA 1130/2560) in accordance with the U.K. Animal (Scientific Procedure) Act, 1986 and the National Institutes of Health guide for the care and use of Laboratory animals (NIH Publications No. 8023, revised 1978).

#### 2.5.2 Anaesthetised rat preparation

Male Sprague-Dawley rats weighing 300-350 g were anaesthetised using Zoletil^®^ (20 mg/kg) and Xylazine^®^ (5 mg/kg) via the intraperitoneal (i.p.) route. Additional anaesthetic was administered throughout the experiment as required. A midline incision was made in the cervical region, and cannulae were inserted into the right jugular vein (for antivenom administration), carotid artery (for measurement of blood pressure and sample collection) and the trachea (for artificial respiration, if required). Arterial blood pressure was recorded using a reusable pressure transducer filled with heparinised saline (25 U/mL). Systemic blood pressure was monitored on a MacLab system (ADInstruments). The rats were kept under a heat lamp during the experiment. At the conclusion of the experiment, the animals were euthanised by an overdose of pentobarbitone (i.v.).

#### 2.5.3 Venom dose optimisation

Preliminary experiments examined the nephrotoxic effects of *D. siamensis* venom via intramuscular (i.m.) doses of 100 µg/kg (*e.g.* 30 µg for 300 g rat), 200 µg/kg (*e.g*. 60 µg for 300 g rat) and 700 µg/kg (*e.g.* 210 µg for 300 g rat) (*n* = 3 per venom dose). Venom was dissolved in 0.9% NaCl and administered i.m. using a 27-gauge needle into the extensor muscles of the right hind limb. Venom doses < 700 µg/kg failed to induce a significant increase in blood urea nitrogen (BUN) and creatinine within 12 hours. Subsequently, the dose of 700 µg/kg (i.m.) was chosen to study the effectiveness of DSAV in subsequent experiments (Supporting information 1, S1).

#### 2.5.4 Determination of D. siamensis monovalent antivenom effectiveness

Where indicated, monovalent *D. siamensis* antivenom (DSAV, Lot No.: WR00117) at two (*i.e*. 0.7 mL for 300 g rat) and three times (*i.e*. 1.05 mL for 300 g rat) the venom/antivenom ratio of the recommended therapeutic dose (*i.e.* 1 mL antivenom per 0.6 mg *D. siamensis* venom) was manually administered via the jugular vein at an infusion rate of 0.25 mL/min over 3-4 min. Control rats were injected with the same volume of normal saline (0.9% sodium chloride, i.v.). Antivenom was administered 15 min prior, or 1 h after, venom administration.

#### 2.5.5 Blood collection for determination of creatinine and blood urea nitrogen (BUN)

At various time points during the animal experiments (0, 3, 6, 9, and 12 h post-injection of venom or 0.9% NaCl), approximately 0.5 mL of blood was taken via the carotid artery and collected in to 1.5 mL Eppendorf tubes. After collection, the samples were centrifuged at 5,500 rpm for 10 min. The supernatant was stored at –20 °C for no longer than 12 h, before determination of creatinine and BUN levels. Creatinine and BUN levels were measured at 37 °C via an automated process using Flex^®^ reagent cartridges and a Dimension^®^ clinical chemistry system supplied by Siemens Healthineers (Germany). Plasma BUN values were measuring using 340 and 383 nm wavelengths by bichromatic rate, whereas plasma creatinine level was measured using 540 and 700 nm wavelengths using bichromatic rate.

#### 2.5.6 Data Analysis and Statistics

Increases in plasma BUN and creatinine were calculated by subtracting the values of the control group from the treatment group, and then presented as mean ± standard error of the mean (SEM). The 95% confidence interval (95% CI) was also calculated. All statistical analyses were performed using GraphPad Prism 6 (GraphPad Software Inc., USA). Multiple comparisons were made using one-way analysis of variance (ANOVA) followed by Bonferroni’s multiple comparison test. Statistical significance was indicated where *P* < 0.05.

## 3. Results

### 3.1 Comparison of the end-point titre of antivenoms against D. siamensis venoms

To compare the immunological binding of *Daboia siamensis* antivenom (DSAV) with other monovalent antivenoms (*i.e. Calloselasma rhodostoma* and *Trimeresurus albolabris* monovalent antivenoms, CRAV and TAAV respectively) and the Hemato Polyvalent antivenom (HPAV) against *D. siamensis* venom, we performed end-point titration ELISA experiments. First, the concentration of each equine F(ab’)_2_ antivenom was standardised to 50 mg/ml before ELISAs were performed with *D. siamensis* venom from three geographical localities: Thailand, Myanmar and Taiwan. Overall, the patterns of immunological binding, as evidenced by an initial plateau and then subsequent decline of OD value (405 nm) after successive antivenom dilutions, was strikingly similar for each of the three venoms tested (Figure 2). The OD readings of the various antivenom/venom combinations at the 1:2,500 dilution provide the most discriminatory comparison and, for clarity, are presented in Table 1. The general trend, including at this dilution, revealed that the antibody-venom binding levels are highest when using the HPAV and DSAV antivenoms, with both displaying considerably higher binding levels to the three different *D. siamensis* venoms than that of the TAAV and CRAV monovalent antivenoms. These results were anticipated, as *D. siamensis* venom is used as an immunogen for both the HPAV (among other venoms) and the DSAV products. While the binding trends are similar, the Taiwanese Russell’s viper venom displayed the lowest binding to both DSAV and HPAV, while venom from Thailand showed the highest binding activity to all of the antivenoms (Figure 2; Table 1).

**Table 1.** Comparison of the immunological binding between the various antivenoms and the three geographical venom variants of *D. siamensis*. The table displays the optical density readings (405 nm) at 1:2,500 dilutions of the antivenoms determined by end-point titration ELISA experiments. Data displayed are means (±SD) of triplicate OD readings (*n* = 3). ** indicates the venom used to raise the antibodies.

| Location of *D. siamensis* venom | Antivenom |  |  |  |
| --- | --- | --- | --- | --- |
|  | Hemato Polyvalent (HPAV) | *D. siamensis* (DSAV) | *C. rhodostoma* (CRAV) | *T. albolabris* (TAAV) |
| Thailand** | 2.19 ± 0.02 | 2.14 ± 0.03 | 1.09 ± 0.04 | 1.37 ± 0.02 |
| Myanmar | 2.13 ± 0.11 | 2.00 ± 0.16 | 0.74 ± 0.04 | 1.05 ± 0.11 |
| Taiwan | 1.86 ± 0.12 | 1.85 ± 0.03 | 0.67 ± 0.04 | 0.91 ± 0.02 |

**Figure 2.**
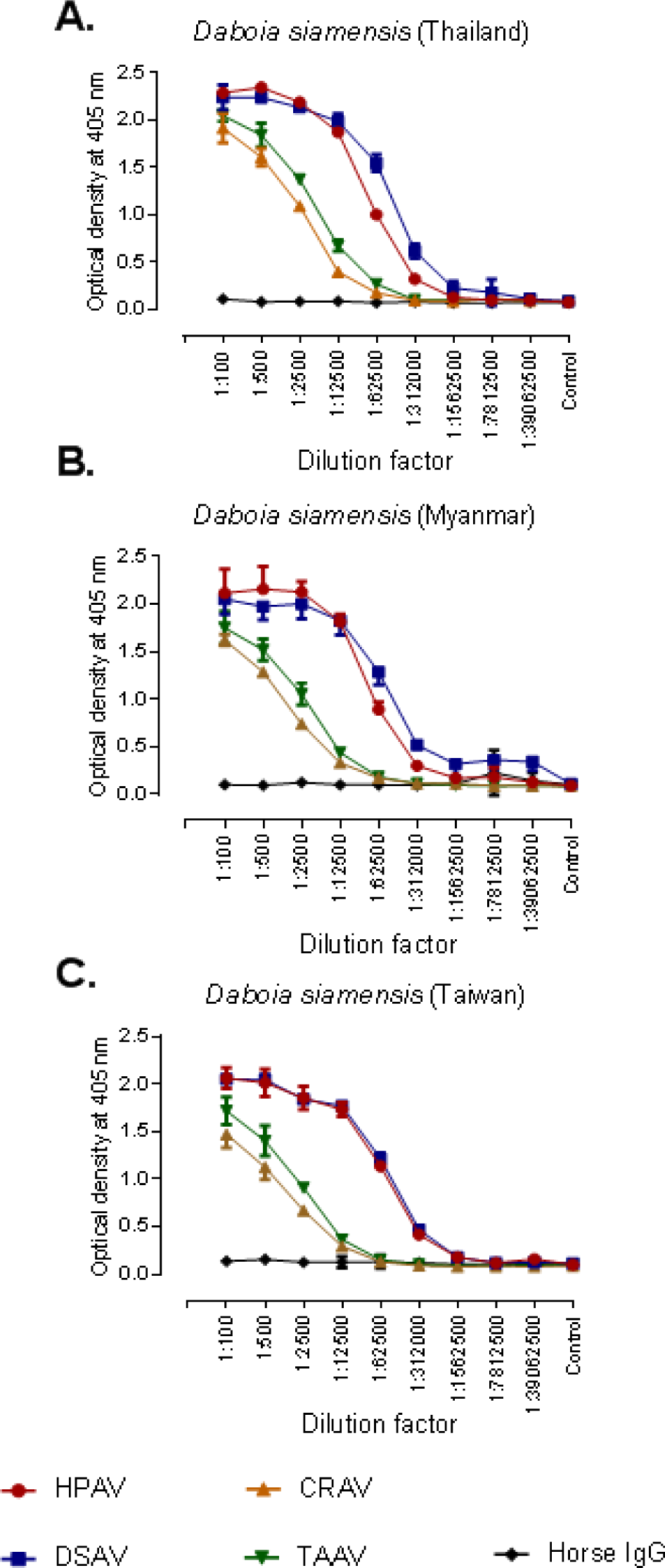
Hemato Polyvalent (HPAV) and *D. siamensis* monovalent (DSAV) antivenoms show extensive and comparable immunological binding to three geographical venom variants of *D. siamensis*. Line graphs show the immunological cross-reactivity of four commercial antivenoms from the Thai Red Cross Society and the negative control (normal horse IgG) against *D. siamensis* venoms from Thailand (A), Myanmar (B) and Taiwan (C) as determined by end-point titration ELISA. Dilution factors are displayed on the x-axis and all antivenoms were adjusted to 50 mg/ml prior to dilution. The control (on the x-axis) represents no venom. Data points represent means of triplicate measurements, and error bars represent SEM.

### 3.2 Comparison of the avidity of antivenoms against D. siamensis venoms

To determine the strength of venom-antivenom antibody binding, we performed avidity ELISAs using a chaotrope to disrupt protein-protein interactions (ammonium thiocyanate, NH_4_SCN). The assay was performed by exposing the same four antivenoms and three *D. siamensis* venoms to increasing concentrations of NH_4_SCN, before reading binding levels by OD (405 nm). Consistent with our findings from the EPT ELISA assay, the venom interactions with HPAV, closely followed by DSAV, were least affected by the presence of the chaotrope, as evidence by the lowest percentage reduction in OD after 4M NH_4_SCN treatment against each of the three Russell’s viper venoms (Figure 3, Table 2). The avidity of both these antivenoms against all three *D. siamensis* venoms was considerably stronger than that observed with CRAV and TAAV (Figure 3, Table 2). However, in contrast to the EPT ELISA, the strength of binding varied among the geographical variants of *D. siamensis* tested, with the greatest avidity detected with the Thai venom used as an immunogen, and lowest avidity observed with the Taiwanese and the Myanmar venoms (Figure 3, Table 2).

**Table 2.**
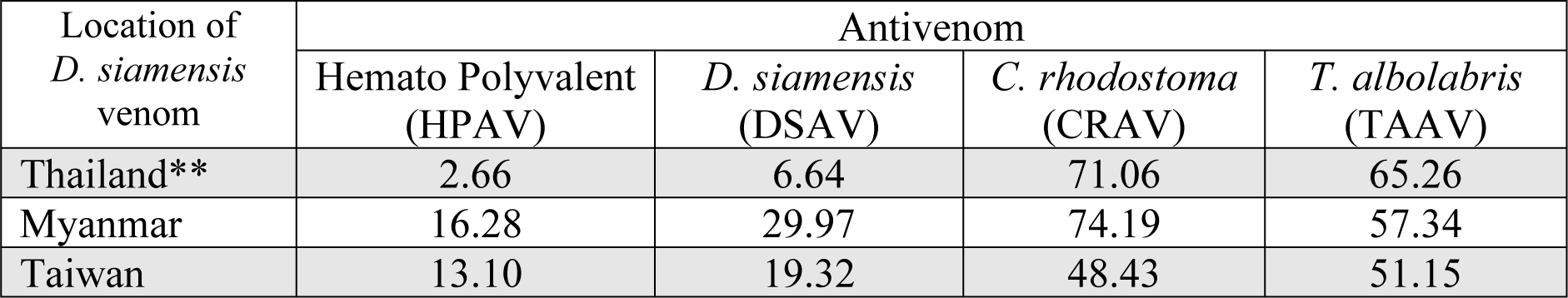
Comparisons of the avidity between the various antivenoms and the three geographical venom variants of *D. siamensis*. The table displays the percentage reduction in optical density (405 nm) readings after the addition of 4M NH_4_SCN as a chaotrope, as determined by avidity ELISA experiments. ** indicates the venom used to raise the antibodies.

**Figure 3.**
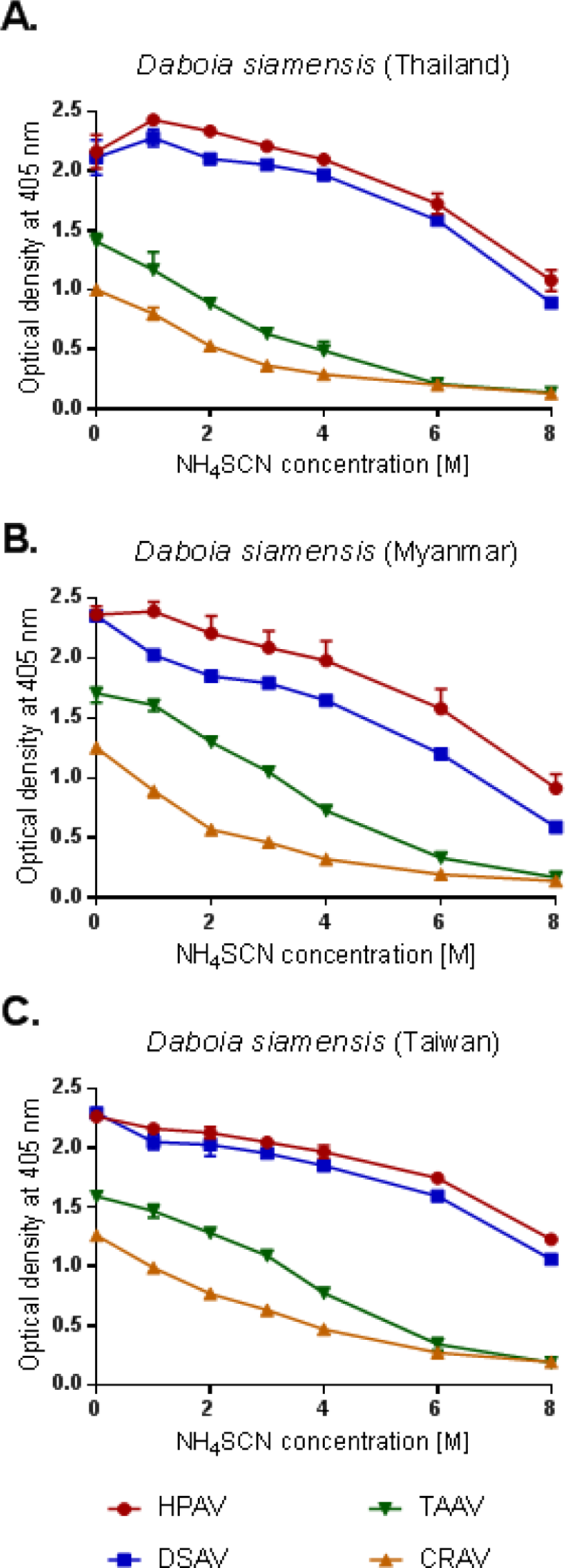
Hemato Polyvalent (HPAV) and *D. siamensis* monovalent (DSAV) antivenoms show high avidity to the toxins found in three geographical venom variants of *D. siamensis*. The avidity of four commercial antivenoms from Thai Red Cross Society against *D. siamensis* venoms from Thailand (A), Myanmar (B) and Taiwan (C) as determined by avidity ELISA. All antivenoms were standardised to 50 mg/ml and used at 1:1,000 dilutions before incubation with NH_4_SCN at increasing molar concentration for 15 minutes. Data points represent means of triplicate measurements, and error bars represent SEM.

### 3.3 Visualising the specificity of antivenoms against D. siamensis venoms

To visualise the specificity of the various antivenoms against the venoms of *D. siamensis* from Thailand, Myanmar and Taiwan, we performed SDS-PAGE gel electrophoresis and western blotting experiments. The venoms (9 µg) were first resolved in a 15% SDS gel under reducing conditions. Our analysis shows that the three venoms have broadly similar venom profiles, with a variety of proteins detected across a large molecular weight range in each sample (Figure 4A). There is, however, a degree of variation in the toxic constituents observed, both in terms of the intensity of shared venom components, and the unique presence of protein bands in some instances (Figure 4A). Notably, a high degree of similarity was observed between the *D. siamensis* venoms from Thailand and Taiwan, whereas the venom from Myanmar exhibited a distinct protein pattern at 12-13 kDa, which is consistent with prior analyses of Myanmar Russell’s viper venom [28]. Nonetheless, western blotting experiments with HPAV and the DSAV against each of the three *D. siamensis* venoms revealed extensive immunological recognition (Figure 4B and 4C, respectively). In each case, the vast majority of venom components observed in the SDS-PAGE profiles were recognised by the antibodies of the two antivenoms with high intensity, and little variation was observed between the two antivenoms (Figure 4B and 4C). In contrast, the CRAV and TAAV monovalent antivenoms displayed almost a complete absence of immunological recognition to the various *D. siamensis* venoms tested (Figure 4D and 4E, respectively).

**Figure 4.**
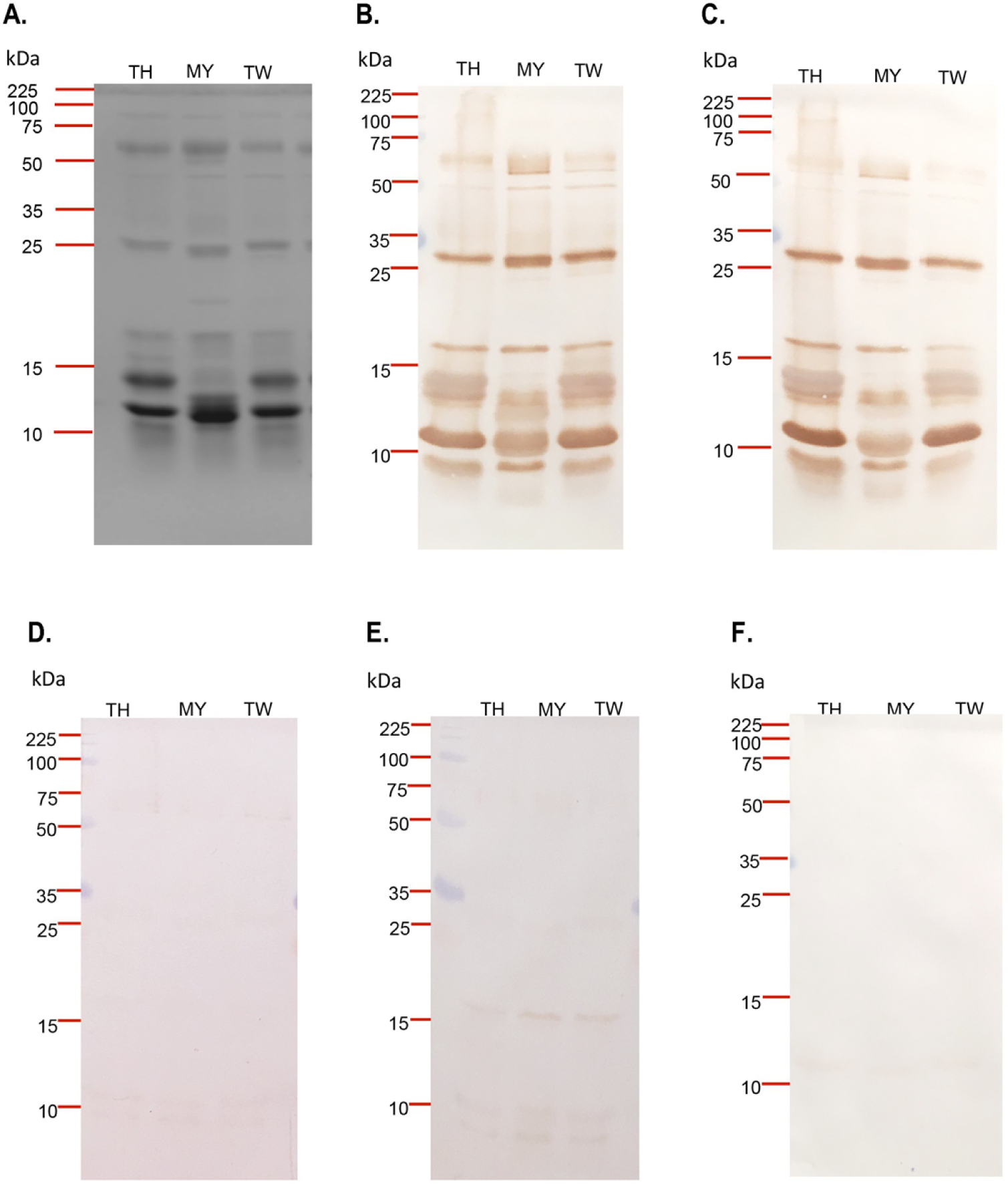
The protein profiles of the three *D. siamensis* venoms and their immunorecognition when probed with antivenoms from the Thai Red Cross Society. (A) SDS-PAGE analysis of *D. siamensis* venoms from Thailand (TH), Myanmar (MY) and Taiwan (TW). Western blotting experiments performed with the three *D. siamensis* venoms and (B) the Hemato Polyvalent antivenom (HPAV), (C) the *D. siamensis* monovalent antivenom (DSAV), (D) the *C. rhodostoma* monovalent antivenom (CRAV), (E) the *T. albolabris* monovalent antivenom (TAAV) and (F) the negative control (normal horse IgG).

### 3.4 Quantifying the coagulopathic venom effects and their neutralization by antivenom

We next quantified the coagulopathic effect of the Thai *D. siamensis* venom (100 ng) using a small scale plasma coagulation assay, which revealed rapid and potent coagulation, consistent with previous studies using *D. russelii* venom [27]. Following the addition of DSAV at 1×, 2× and 3× the scaled recommended therapeutic dose, we observed significant inhibition of coagulation with each antivenom treatment (*p* < 0.05 *vs* venom only control) (Supporting Information S2). We therefore used the 1× recommended therapeutic dose of DSAV (3.6 µg/well) as a potentially discriminatory dose to compare the relative neutralizing capability of the three other antivenoms (HPAV 9.2 µg/well, CRAV 1.7 µg/well and TAAV 3.6 µg/well) against the Thai *D. siamensis* venom. HPAV exhibited significant inhibition of coagulopathic venom activity, in a manner highly comparable with DSAV. Consistent with the lower levels of immunological binding observed in our earlier experiments, the CRAV and TAAV monovalent failed to inhibit the coagulopathic activity of Thai Russell’s viper venom (Supporting Information, S2).

Finally, we assessed the ability of the two antivenoms (DSAV and HPAV) exhibiting neutralizing potential against the Thai *D. siamensis* venom, to neutralize the coagulopathic venom effects of *D. siamensis* venoms from Myanmar and Taiwan. Both venoms (100 ng) caused rapid clotting activity comparable with that of the Thai venom (Figure 5). However, both the DSAV and HPAV antivenoms at 1× the scaled recommended therapeutic dose (*i.e.* 1 mL per 0.6 mg of *D. siamensis* venom) prevented the rapid coagulation induced by *D. siamensis* venom from Thailand, Myanmar and Taiwan (*n* = 3, *P*< 0.05, one-way ANOVA, followed by Bonferroni’s *t*-test, Figure 5).

**Figure 5.**
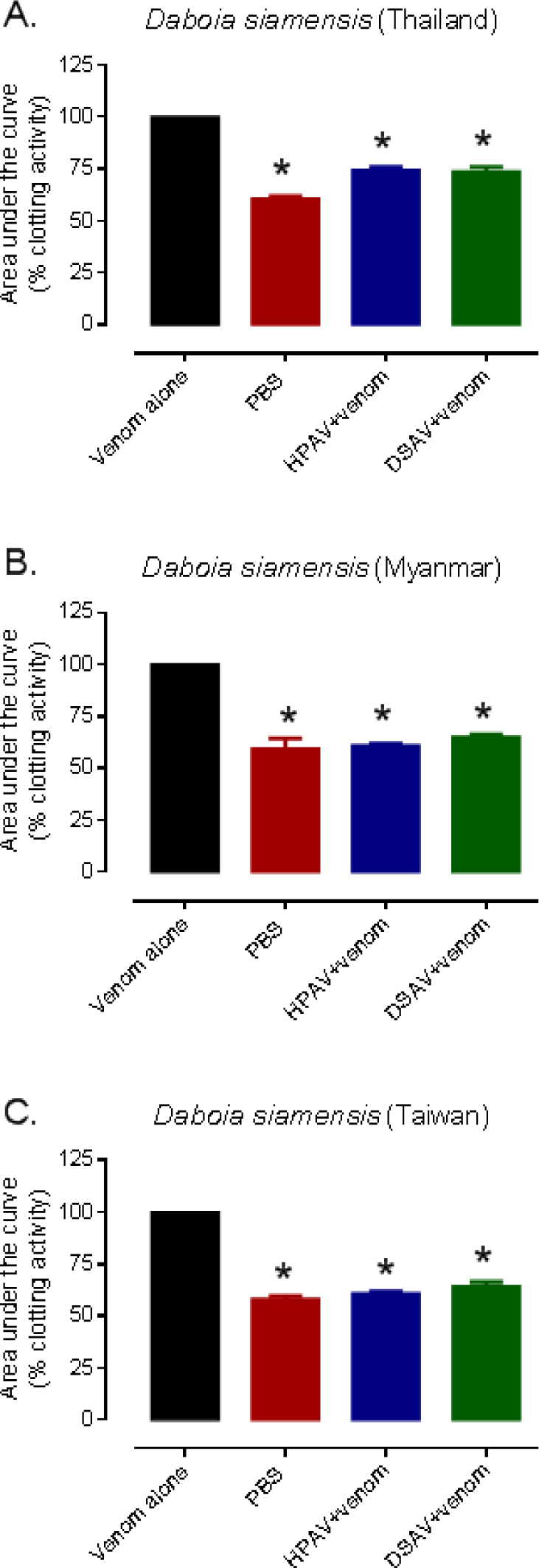
The procoagulant activity of the three different *D. siamensis* venoms and their neutralization by the DSAV and HPAV antivenoms. (A) Thailand, (B) Myanmar, and (C) Taiwan. The antivenoms were tested at the manufacturer’s recommended therapeutic dose. The data represents kinetic profiles of clotting from the plasma coagulation assay displayed as mean areas under the curve from triplicate measurements, transformed into percentage of the venom only control, and error bars represent SEM. * *P* < 0.05, compared to *D. siamensis* venom alone (one-way ANOVA, followed by Bonferroni *t*-test).

### 3.5 The effectiveness of DSAV on Russell’s viper-induced nephrotoxicity

A significant increase in plasma BUN levels were observed following the administration of *D. siamensis* venom (700 µg/kg) via the intramuscular (i.m.) route into the anaesthetised rat, when compared to the control group (Supporting information, S1). Time course sampling (every three hours) revealed that BUN increased at each time point up to the end of the experiment (12 hrs, Figure 6A). The intravenous administration of DSAV (i.v.) at 3× the scaled recommended therapeutic dose (*i.e.*, 1 mL per 0.6 mg of *D. siamensis* venom) prior to the injection of venom resulted in a significant reduction in plasma BUN levels compared to the venom only controls (*n*=4-5, *P* < 0.05) (Figure 6A). However, no significant reduction in BUN levels was observed with a reduced therapeutic dose of 2x that recommended. The administration of antivenom 1 h after the i.m. administration of venom also did not significantly decrease plasma BUN levels compared to the administration of venom alone (*n* = 4-5, *P* < 0.05, one-way ANOVA, followed by Bonferroni’s *t*-test, Figure 6B).

**Figure 6.**
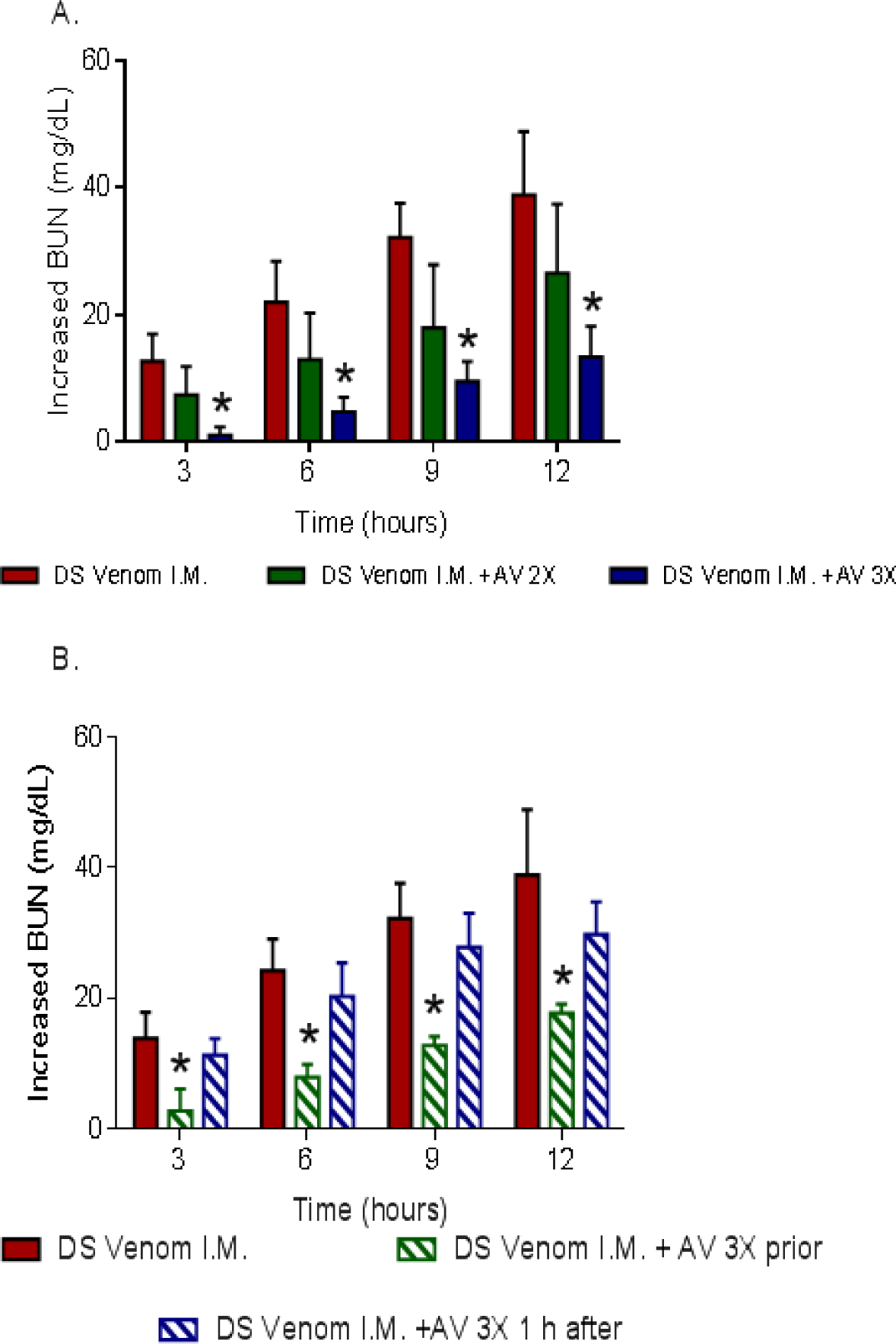
High doses of *D. siamensis* monovalent (DSAV) antivenom are required to abrogate increased plasma BUN levels caused by the administration of Thai *D. siamensis* venom. (A) The graphs show increases in the blood urea nitrogen (BUN) concentrations of rats administered with (i) *D. siamensis* venom (700 µg/kg, i.m.), and (ii) venom alongside the pre-administration of DSAV at two times the recommended therapeutic dose and (iii) venom alongside the pre-administration of DSAV at three times the recommended therapeutic dose. (B) Prior administration of DSAV at three times the recommended therapeutic dose significantly prevented the increase plasma BUN compared with antivenom given 1 h after venom. Data is displayed for BUN of rats administered with (i) *D. siamensis* venom (700 µg/kg, i.m.), (ii) venom alongside the pre-administration of DSAV at three times the recommended therapeutic dose, and (iii) venom and antivenom (3x recommended dose) 1 hr after venom administration. The data displayed is presented as increased levels compared to the control (normal saline, *n*=4-5) and represent mean measurements (*n*=4-5), with error bars representing SEM. * *P* < 0.05, compared to *D. siamensis* venom alone (one-way ANOVA, followed by Bonferroni *t*-test).

In addition to BUN, the intramuscular administration of *D. siamensis* venom (700 µg/kg) also resulted in significant increases in plasma creatinine levels compared to the control group (Figure 7A and B). Creatinine levels also increased over time and were significantly reduced when DSAV at 3× the recommended therapeutic dose (*n* = 4-5, *P* < 0.05) was intravenously administration prior to the injection of venom, but no significant effect was observed when 2× the recommended dose was administered (Figure 7A). However, in contrast with BUN, the administration of antivenom (i.v., infusion; 3× recommended titre) 1 h after the i.m. administration of venom caused a significant decrease in plasma creatinine compared to the administration of venom alone (*n* = 4-5, *P* < 0.05, one-way ANOVA, followed by Bonferroni’s *t-*test, Figure 7B).

**Figure 7.**
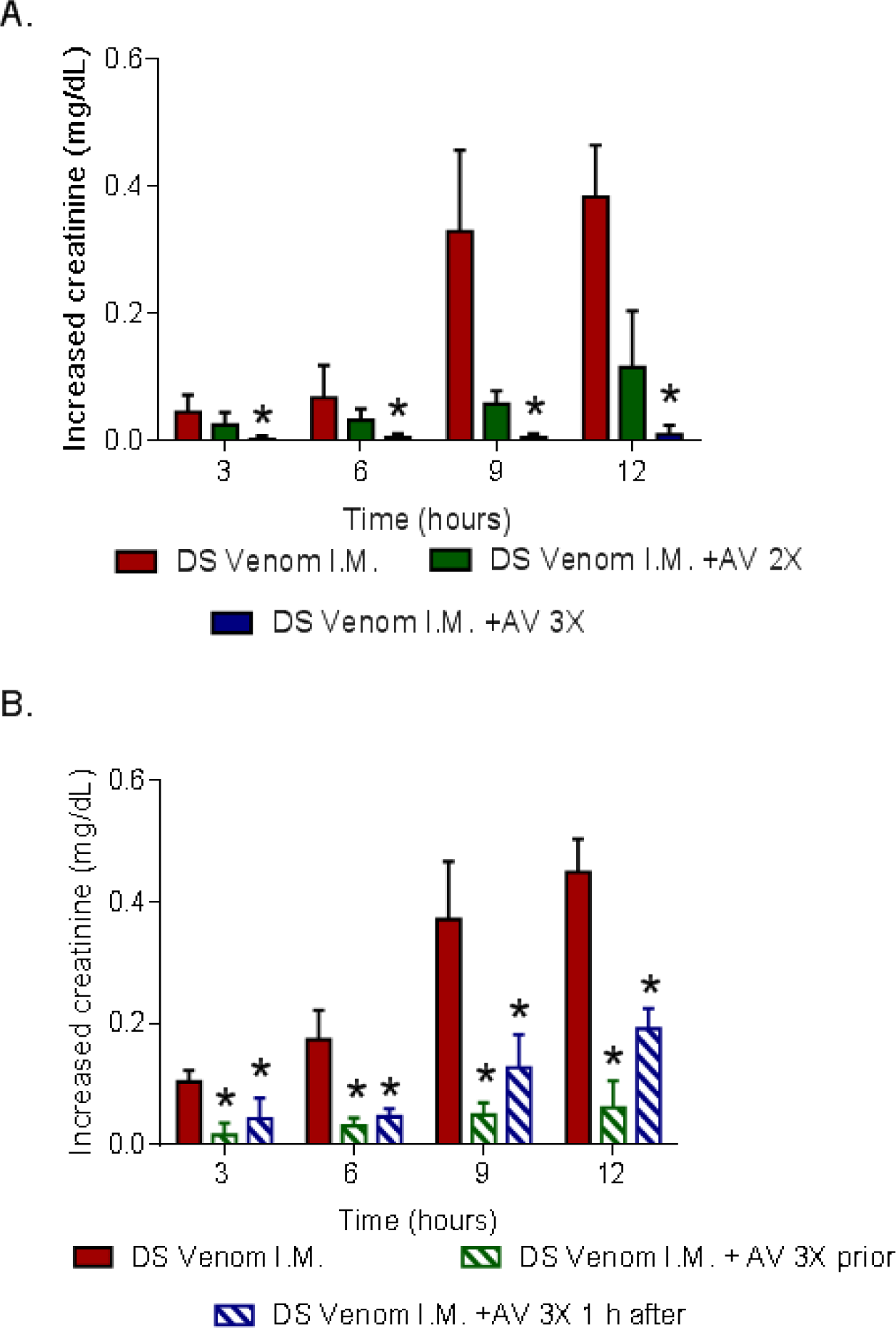
High doses of *D. siamensis* monovalent (DSAV) antivenom are required to abrogate increased plasma creatinine levels caused by the administration of Thai *D. siamensis* venom. (A) Plasma creatinine concentrations of rats administered with (i) *D. siamensis* venom (700 µg/kg, i.m.), and (ii) venom alongside the pre-administration of DSAV at two times the recommended therapeutic dose and (iii) venom alongside the pre-administration of DSAV at three times the recommended therapeutic dose. (B) Delayed administration of DSAV still results in significantly reduced plasma creatinine levels induced by *D. siamensis* venom *in vivo*. The graphs show plasma creatinine concentrations of rats administered with (i) *D. siamensis* venom (700 µg/kg, i.m.), (ii) venom alongside the pre-administration of DSAV at three times the recommended therapeutic dose, and (iii) venom and antivenom (3x recommended dose) 1 hr after venom administration. The data displayed is presented as increased levels compared to the control (normal saline, *n*=4-5) and represents mean measurements (*n*=4-5), with error bars representing SEM. * *P* < 0.05, compared to *D. siamensis* venom alone (one-way ANOVA, followed by Bonferroni *t*-test).

## 4. Discussion

Snakes of the genus *Daboia* (Russell’s vipers; *D. siamensis* and *D. russelii*) are widely distributed across Asia and bites by these species cause thousands of fatalities each year. The mainstay of treatment for Russell’s viper envenoming is the administration of polyclonal antibodies, known as monovalent or polyvalent antivenoms. However, the treatment of systemic envenoming caused by *D. siamensis* has long been problematic in many Asian countries, due to challenges related to the access of antivenom and the dosing regimen used to effect cure. For example, in southern parts of mainland China, where envenoming by *D. siamensis* poses a substantial health problem, access to monovalent antivenoms is very limited, which has resulted in the use of non-specific or species-inappropriate antivenoms, leading to reports of treatment failures and mortality [17]. Herein, we examined the effectiveness of the Thai monovalent *D. siamensis* (DSAV) and Hemato Polyvalent (HPAV) antivenoms against three different geographical variants of *D. siamensis* using *in vitro* biochemical and immunological assays. We also investigated the preclinical efficacy of the DSAV antivenom against the nephrotoxic effects of *D. siamensis* venom.

We first used a range of immunological assays to assess the amount of binding, strength of binding and specificity of antivenom antibodies against *D. siamensis* venoms from Thailand, Myanmar and Taiwan. Both end point titration and avidity ELISA experiments demonstrated substantial cross-reactivity between all three *D. siamensis* venoms and the DSAV and HPAV antivenoms, and very little cross-reactivity with the control antivenoms used (CRAV and TAAV; neither of these products use *D. siamensis* as an immunogen). The EPT ELISA showed that *D. siamensis* venoms from Myanmar and Taiwan are well recognised by these two commercial antivenoms, with binding levels highly comparable to those observed with the Thai venom (Figure 2), which was used for immunization during antivenom production. The avidity ELISA was more discriminatory, with the strength of antibody-venom protein binding greatest for both antivenoms against the Thai *D. siamensis* venom (Figure 3). These results are in line with a previous study, which showed that the Thai DSAV antivenom exhibits immunoreactivity to *D. siamensis* venoms from Taiwan and Guanxi, South China, but to a lesser extent than the binding observed with Chinese monovalent antivenom [17].

Prior proteomic studies have demonstrated that *D. siamensis* venom from Myanmar contains at least six major protein families; serine proteinases, metalloproteinases, PLA_2_, L-amino acid oxidases, vascular endothelial growth factors and C-type lectin-like proteins [9]. In our SDS-PAGE analysis, we find that *D. siamensis* venom from Myanmar displayed high intensity protein bands at around 10-15 kDa, which differed from the highly comparable venom protein profiles of the Taiwanese and Thai *D. siamensis* venoms (Figure 4A). However, western blotting experiments showed that both the DSAV and HPAV antivenoms recognise the vast majority of venom proteins present in these venoms, despite the element of venom variation present in the Myanmar geographical variant (Figure 4B and C). The exception to this is perhaps proteins observed in the 50-100 kDa molecular weight range, where lower binding between both antivenoms and the venoms was observed, with those from Myanmar and Taiwan exhibiting the lowest cross-reactivity. While immunological assays alone cannot be used to define the likely preclinical efficacy of an antivenom [26, 29], strong immunological characteristics are an essential prerequisite for venom neutralization *in vivo*. Thus, our findings from ELISA and immunoblotting experiments suggest that the Thai DSAV and HPAV antivenoms may neutralise *D. siamensis* venom from different parts of its range, and thus may be a useful clinical tool across Southeast Asia. However, this assertion needs to next be validated in future studies using preclinical models of antivenom efficacy.

Envenoming by snakes of the genus *Daboia* manifest in a variety of clinical outcomes. For example, in Sri Lanka, some bites by *D. russelii russelii* have been reported to cause neurotoxicity characterized by flaccid paralysis, myotoxicity associated with skeletal muscle breakdown, and coagulopathy [30, 31]. In the case of *D. siamensis*, two of the most severe and common clinical outcomes observed following envenoming by this species are systemic coagulopathy and acute renal failure [7, 32]. Unfortunately, such signs are common when antivenom therapy is delayed or absent, and in a prior study resulted in over 70% of systemically envenomed Taiwanese victims presenting with thrombocytopenia, hemolysis and acute renal failure [33]. Russell’s viper venom is thought to cause systemic coagulopathy via procoagulant toxins (e.g. RVV-X and RVV-V) potently activating the clotting factors Factor X and Factor V [34]. Continual activation of the blood coagulation cascade results in the depletion of clotting factors, most notably fibrinogen, and results in an incoagulable blood syndrome known as VICC [35, 36]. The presence of VICC makes victims highly vulnerable to suffering from severe haemorrhages, which can be lethal, particularly if bleeds occur intracranially [37].

To assess the ability of the Thai antivenoms to neutralise the coagulopathic effects of *D. siamensis* venom, we used a plasma coagulation assay previously validated using Russell’s viper venom [27]. All three *D. siamensis* venoms exerted strong procoagulant effects in a comparable manner, but this venom activity was effectively neutralised by the DSAV, and to a lesser extent by the HPAV, at the scaled recommended therapeutic dose (*i.e.* 1 mL antivenom per 0.6 mg of *D. siamensis* venom). We found no significant differences between the neutralising activity of either of these antivenoms against the three different venoms. In contrast, neither the CRAV or the TAAV showed any neutralising activity against Thai *D. siamensis* venom-induced coagulopathy, which is consistent with these venoms being absent from the immunogen mixture,and supports the hypothesis that different venomous snakes cause coagulopathy via different mechanisms [38]. Overall, these findings support the notion that the extensive immunological cross-reactivity observed among the DSAV and HPAV antivenoms and *D. siamensis* venoms translates into neutralisation of venom function, at least in the context of coagulopathic toxins.

Nephrotoxicity is an important complication diagnosed following envenomings by a number of hemotoxic and myotoxic snake species, such as *D. siamensis* and certain sea snakes (subfamily *Hydrophiinae*) [8]. Envenoming by *D. siamensis* has previously been described to cause a number of pathological renal changes including proteinuria, haematuria, rhabdomyolysis and acute renal failure [8]. Acute renal failure has been indirectly linked to other systemic pathologies caused by *D. siamensis* venom, such as intravascular haemolysis, VICC and glomerulonephritis, while direct nephrotoxic activity has also been reported as a cause of renal failure [8]. Prior *in vivo* experiments, which monitored renal hemodynamics in anaesthetised dogs, showed that purified PLA_2_ and SVMP toxins from *D. siamensis* venom played an important role in causing renal vascular changes [20]. Furthermore, rapid increases in plasma BUN and creatinine levels appear to be useful markers for the diagnosis of Russell’s viper venom-induced acute renal failure [32]. In particular, elevation in plasma creatinine appears to be a significant biomarker indicating nephrotoxicity induced by snake envenomation. For example, a number of studies have shown that changes in plasma creatinine following envenomation by *Pseudechis australis* (mulga snake) or *Crotalus durissus* (neotropical rattlesnake) are associated with acute renal failure in both animals and humans [39–41].

It remains unclear how effective antivenom therapy is at preventing nephrotoxicity caused by *D. siamensis* venom. A previous preclinical study using experimentally envenomed mice indicated that the administration of HPAV 10 minutes prior to venom delivery effectively inhibited hematuria and proteinuria-induced by *D. siamensis* venoms from Thailand and Myanmar [23]. In this study, we used an anaesthetised rat model, and demonstrated that the intramuscular delivery of *D. siamensis* venom (*i.e.* 700 µg/kg) results in marked increases in both BUN and creatinine. We found progressive increases in renal dysfunction up to the end of our experiments (12 h post-venom administration), with both plasma BUN and creatinine levels increasing at every 3 h sampling point. DSAV administered prior to venom, or 1 h after venom delivery, significantly reduced increases in plasma creatinine concentration, but only had a significant effect on reducing BUN levels when the antivenom was administered prior to the venom. These findings are in general agreement with clinical observations from *D. siamensis* envenoming, where the earlier administration of antivenom prevented renal failure, whereas late treatment (>3 h) did not inhibit renal dysfunction, as determined by increases in serum-creatinine levels [32, 33]. However, further studies are required to investigate whether some of the nephrotoxic effects of *D. siamensis* venom are not effectively inhibit by antivenom, as in the case of the BUN levels monitored here. Moreover, a relatively high volume (*i.e*., three times the recommended scaled therapeutic dose) of the DSAV monovalent antivenom was required to reduce plasma creatinine and BUN levels herein, with therapeutic doses twice that recommended found to have no significant effect. Consequently, the administration of high initial doses of antivenom, with repeated doses subsequently, has been clinically recommended in the presence of rebound antigenemia and recurrent toxicity [42]. Our preclinical findings demonstrate that higher therapeutic doses of antivenom than currently recommended may be required to prevent severe renal toxicity, and this seems likely to be particularly relevant when patients present to hospital in a delayed manner.

### Conclusion

In this study, we demonstrate that the Thai monovalent and polyvalent antivenoms, *i.e.* DSAV and HPAV, exhibit extensive immunological binding and *in vitro* neutralizing effects against *D. siamensis* venoms from Myanmar and Taiwan, and at comparable levels to the Thai venom used to make the antivenom. These findings suggest that these antivenoms may be useful therapeutic agents across much of Southeast Asia, particularly in the event that local antivenom supply is insufficient for the needs of the many snakebite victims. We also demonstrate in a preclinical model that the early administration of high doses of DSAV antivenom may be effective at preventing acute kidney injury, although further work needs to be undertaken to better understand the nephrotoxic effect of *D. siamensis* venom and the disparity between its effect on reducing the BUN and creatine levels described herein. To this end, further work is needed to assess the neutralising effect of antivenom against nephrotoxicity caused by purified toxins from Russell’s viper venoms, to better understand venom-induced acute renal failure and the efficacy of snakebite therapies against this important pathological syndrome.

## Abbreviations

ELISA: enzyme-link-immunosorbent assay
BUN: blood urea nitrogen
NH_4_SCN: ammonium thiocyanate
DIC: disseminated intravascular coagulation
VICC: venom-induced consumption coagulopathy
PBS: phosphate-buffered saline
PLA_2_: phospholipase A_2_ enzymes
SVMP: snake venom metalloproteinase
HPAV: Hemato Polyvalent Snake antivenom
DSAV: monovalent antivenom for *Daboia siamensis*
CRAV: monovalent antivenom for *Calloselasma rhodostoma*
TAAV: monovalent antivenom for *Trimeresurus albolabris*

## Acknowledgments

The authors wish to acknowledge Jaffer Alsolaiss and Laura-Oana Albulescu; Centre for Snakebite Research & Interventions, Liverpool School of Tropical Medicine, UK, for immunological and coagulation assay expertise. The authors also wish to acknowledge the Office of Research Development, Phramongkutklao College of Medicine & Phramongkutklao Hospital (ORD, PCM & PMK, Bangkok, Thailand), Thailand Research Fund (TRF) and British Council. Janeyuth Chaisakul was supported by a Newton Fund Thailand-UK researcher Link award 2017-18 (PDG61W0015), and Nicholas Casewell was supported by a Sir Henry Dale Fellowship (200517/Z/16/Z) jointly funded by the Wellcome Trust and Royal Society.

## Supporting Information Legends

**S1 Fig.** *Daboia siamensis* venom (700 µg/kg, i.m., *n*=3) significantly increases (A) plasma creatinine and (B) BUN levels compared with vehicle control (saline, *n*=3) in an anaesthetised rat model of nephrotoxicity. Data points represent readings from plasma samples collected every 3 hrs. * *P* < 0.05, compared to vehicle control (one-way ANOVA, followed by Bonferroni *t*-test).

**S2 Fig.** The procoagulant activity of *D. siamensis* venom and its neutralization by Thai antivenoms. (A) The neutralizing effect of increasing concentrations of *D. siamensis* monovalent antivenom (DSAV) (1×, 2× and 3× recommended therapeutic dose) on the clotting activity of Thai *D. siamensis* venom. (B) The comparative neutralizing effect of monovalent antivenoms made against *D. siamensis* (DSAV), *C. rhodostoma* (CRAV) and *T. albolabris* (TAAV) venom, and the Hemato Polyvalent antivenom (HPAV), on the procoagulant venom activity of Thai *D. siamensis* venom. The coagulation assay kinetically monitors the clotting of bovine plasma, and the data displayed represents areas under the curve of the resulting kinetic profiles, transformed into percentage of the venom only control. Data points represent the means of triplicate measurements, and error bars represent SEM. * *P* < 0.05, compared to *D. siamensis* venom alone (one-way ANOVA, followed by Bonferroni *t*-test).

**S3 Fig.** The kinetic profiles of procoagulant activity of the three different *D. siamensis* venoms and their neutralization by the *D. siamensis* monosvalent (DSAV) and the Hemato Polyvalent (HPAV) antivenoms. (A) Thailand, (B) Myanmar, and (C) Taiwan. The antivenoms were tested at the recommended therapeutic dose (1x). The data displayed is the kinetic profiles from the plasma coagulation assay and data points represent the means of triplicate measurements, and error bars represent SEM. Normal clotting is indicated by the red line (PBS).

